## Supplemental Figures and Supplemental File Descriptions for "GWAS of three molecular traits highlights core genes and pathways alongside a highly polygenic background"

**Supplementary File 1:** *Independent GWAS hits for urate. CHROM, chromosome number; POS, variant position (hg19); ID, variant identifier; REF, reference genome sequence allele; A1, alternative allele; A1_CT, number of A1 alleles; ALLELE_CT, total alleles; A1_FREQ, frequency of A1 allele; MACH_R2, estimated imputation accuracy (INFO); OBS_CT, number of individuals with non-missing data; BETA, effect size of A1 allele; SE, standard error of A1 allele; T_STAT, t-statistic; P, p-value of association between A1 allele and serum urate levels.*

**Supplementary File 2:** *Independent GWAS hits for IGF-1. CHROM, chromosome number; POS, variant position (hg19); ID, variant identifier; REF, reference genome sequence allele; A1, alternative allele; A1_CT, number of A1 alleles; ALLELE_CT, total alleles; A1_FREQ, frequency of A1 allele; MACH_R2, estimated imputation accuracy (INFO); OBS_CT, number of individuals with non-missing data; BETA, effect size of A1 allele; SE, standard error of A1 allele; T_STAT, t-statistic; P, p-value of association between A1 allele and serum IGF-1 levels.*

**Supplementary File 3:** *Independent GWAS hits for testosterone in males. CHROM, chromosome number; POS, variant position (hg19); ID, variant identifier; REF, reference genome sequence allele; A1, alternative allele; A1_CT, number of A1 alleles; ALLELE_CT, total alleles; A1_FREQ, frequency of A1 allele; MACH_R2, estimated imputation accuracy (INFO); OBS_CT, number of individuals with non-missing data; BETA, effect size of A1 allele; SE, standard error of A1 allele; T_STAT, t-statistic; P, p-value of association between A1 allele and serum testosterone levels in males.*

**Supplementary File 4:** *Independent GWAS hits for testosterone in females. CHROM, chromosome number; POS, variant position (hg19); ID, variant identifier; REF, reference genome sequence allele; A1, alternative allele; A1_CT, number of A1 alleles; ALLELE_CT, total alleles; A1_FREQ, frequency of A1 allele; MACH_R2, estimated imputation accuracy (INFO); OBS_CT, number of individuals with non-missing data; BETA, effect size of A1 allele; SE, standard error of A1 allele; T_STAT, t-statistic; P, p-value of association between A1 allele and serum testosterone levels in females.*

**Supplementary File 5:** *Phenotype-level correlations between luteinizing hormone (LH) and testosterone in females and males. Magnitude of correlation and sample sizes are both higher using the XM0lv luteinizing hormone code, but results are consistent across codes.*

**Supplementary File 6:** *Female-specific association with circulating testosterone levels at the FSHB locus. We observe an association between the previously discovered rs11031006 [125, 109] and serum testosterone levels in females. This association was reproduced in the non-British White individuals in UK Biobank. All effects are at rs11031006 with respect to dosage of the A allele.*

**Supplementary File 7:** *Curated core pathways that showed enrichment for each trait GWAS, as discovered by MAGMA[111] at 5% FDR.*

**Supplementary File 8:** *Pathways representing core genes for serum urate biology. Pathway, which class of genes; Gene name, name of gene included in the given pathway.*

**Supplementary File 9:** *Pathways representing core genes for serum IGF-1 biology. Pathway, which class of genes; Gene name, name of gene included in the given pathway.*

**Supplementary File 10:** *Pathways representing core genes for serum testosterone biology. Pathway, which class of genes; Gene name, name of gene included in the given pathway.*

**Supplementary File 11:** *SNP heritabilities and level of population stratification as estimated by LD Score regression [57] using the full set of baseline and cell type specific annotations for each biomarker trait, with height as a baseline. The lower estimates of LD Score regression SNP-based heritability relative to HESS are expected [112]. Best $h^2$ intercept refers to the intercept of the inflation for the best-fitting simulation results in the $h^2$-derived causal SNP estimates (Methods).*

**Supplementary File 12:** Estimates of SNP heritability and fraction of causal variants from GENESIS [22]. $K$, number of mixture components used in the fit of effect sizes. Half sample, 50% downsample of individuals in GWAS to mimic the sex-specific traits. * failed to converge and terminated after a single iteration.

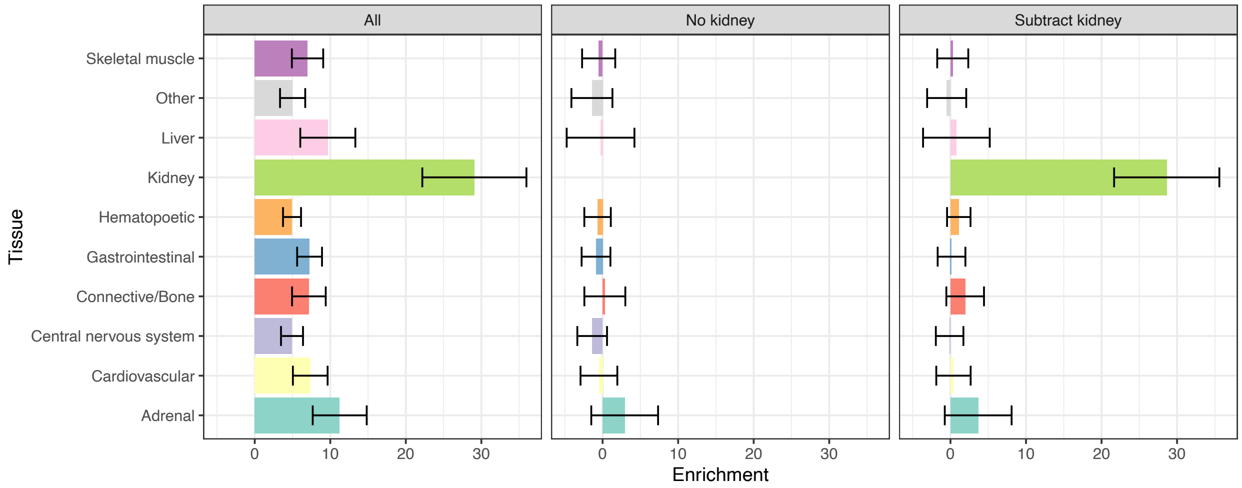


**Figure 1-figure supplement 1:** *Estimates of serum urate SNP-based heritability within cell and tissue group annotations using LD Score regression [57]. Conditions represent naive multiple regression against all cell types simultaneously (left), multiple regression of non-kidney cell types after removing regions overlapping with kidney (middle), and multiple regression of non-kidney cell types excluding kidney regions along with the original kidney annotation (right). 95% confidence intervals are shown. The latter two results suggest that almost none of the urate SNP-based heritability lies specifically within any non-kidney cell type.*

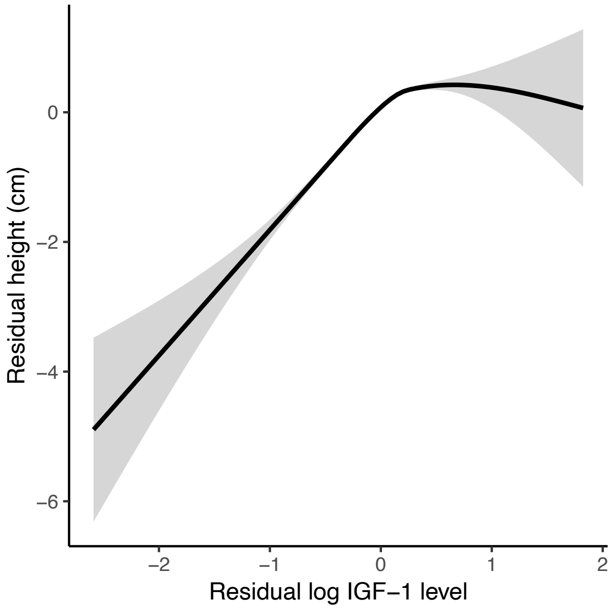


**Figure 3-figure supplement 1:** *Covariate-adjusted IGF-1 levels are significantly associated with covariate-adjusted height in UK Biobank.*

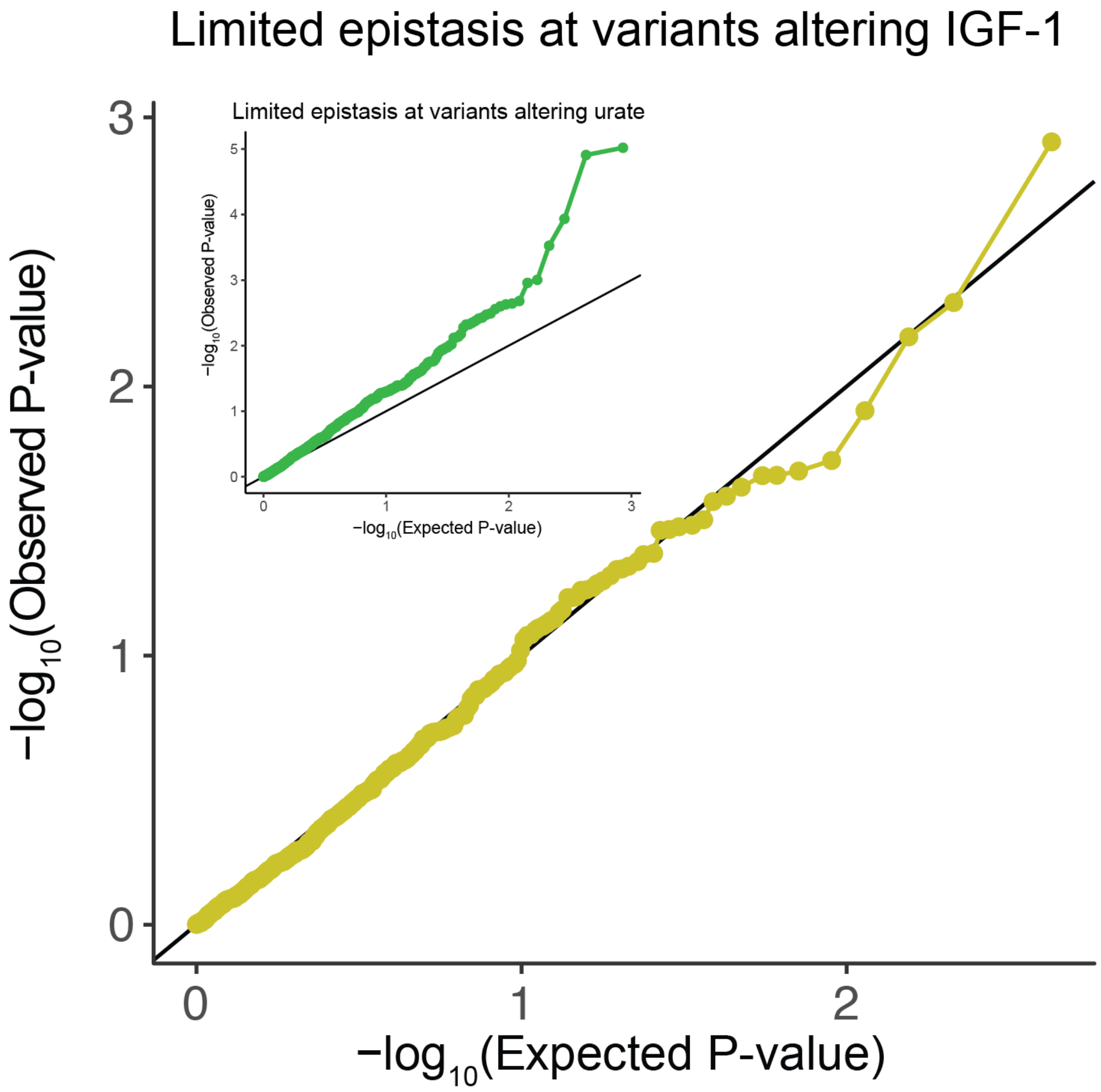


**Figure 3-figure supplement 2:** *QQ-plot testing for epistasis plots all pairs of lead variants with $p < 1e-20$ for IGF-1 levels (Methods). Inset is the corresponding plot for urate levels.*

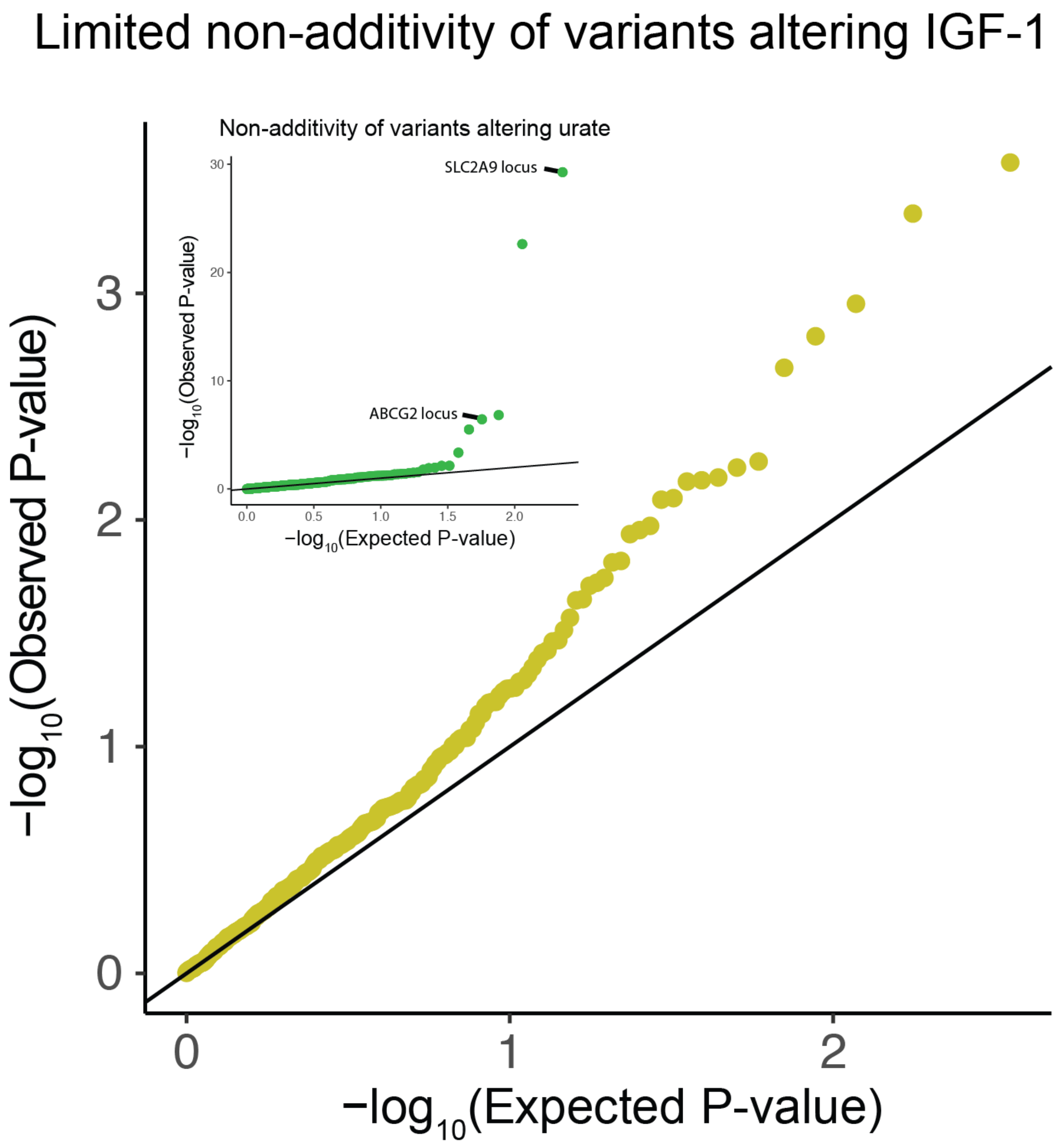


**Figure 3-figure supplement 3:** *QQ-plot testing for non-additivity at IGF-1 associated SNPs. All lead variants with* $p < 5e - 8$ *passing quality control were tested for departures from an additive model (Methods). Inset is the same analysis run on associations with serum urate levels.*

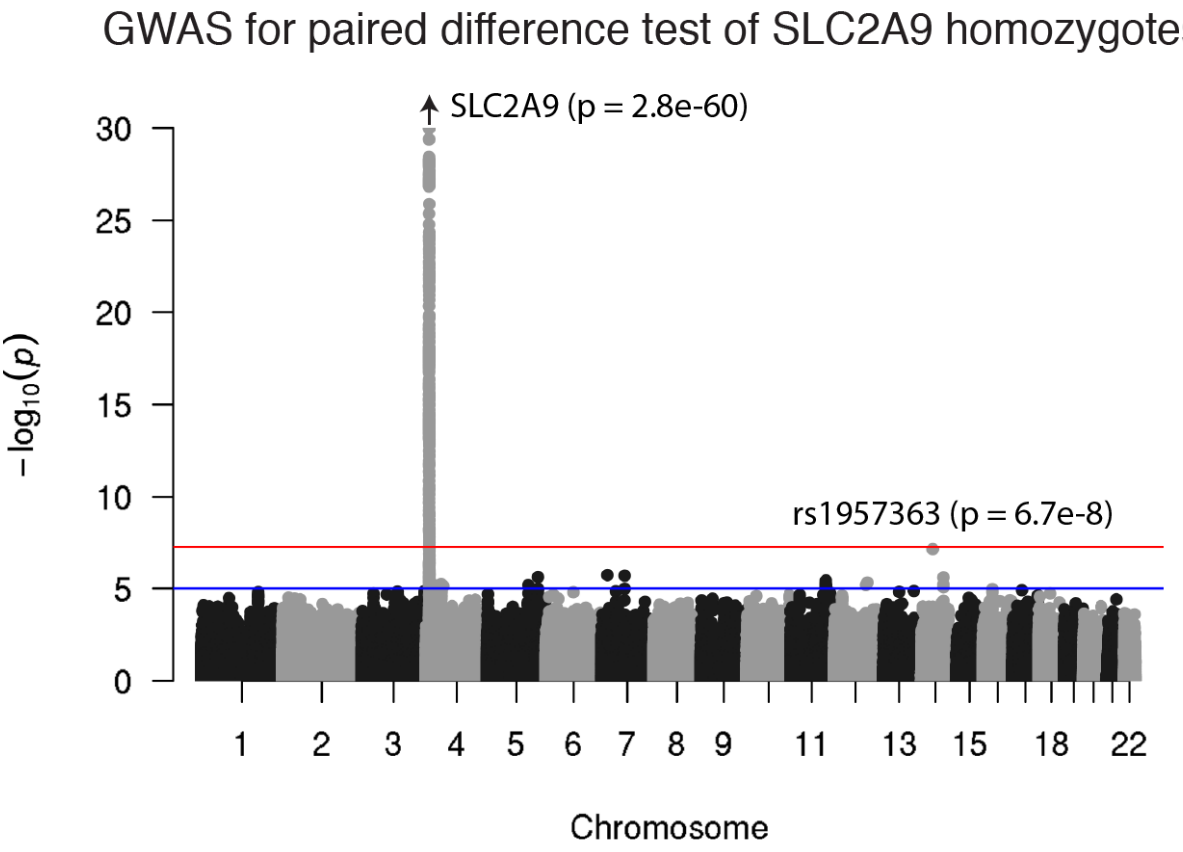


**Figure 3-figure supplement 4:** *A genome wide association study for paired differences in effect size by SLC2A9 genotype. Other than at the SLC2A9 locus, there are no genome-wide significant differences.*

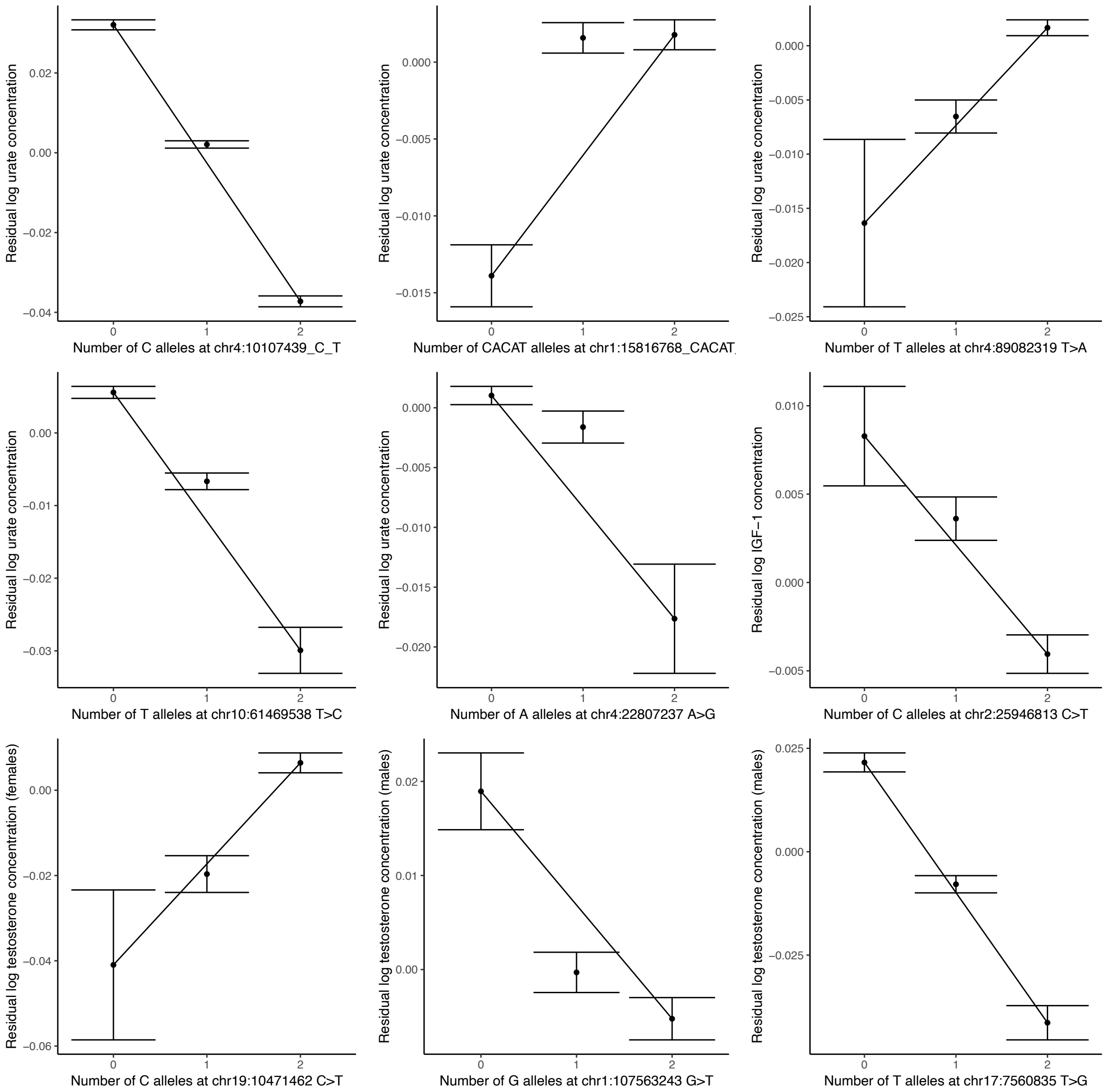


**Figure 3-figure supplement 5:** *Non-additivity in serum urate concentrations at chr4:10107439 C>T, chr1:15816768 CACAT>C, chr4:89082319 T>A, chr4:22807237 A>G, and chr10:61469538 T>A; at chr2:25946813 C>T in IGF-1; and in female testosterone at chr19:10471462 C>T and male testosterone at chr1:107563243 G>T and chr17:7560835 T>G. 95% confidence intervals shown, and lines are drawn between homozygotes. Only chr1:15816768 CACAT>C, chr1:107563243 G>T, and chr4:22807237 A>G show substantial departure from additivity (revealing a recessive effect in all three cases).*

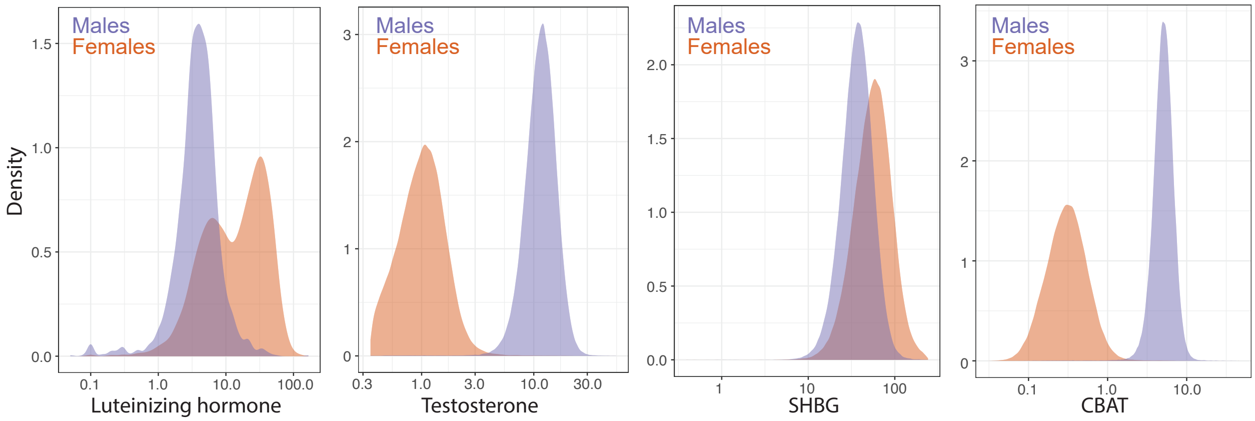


**Figure 5-figure supplement 1:** *Distributions of female and male luteinizing hormone, testosterone, sex hormone binding globulin (SHBG), and calculated bioavailable testosterone (CBAT) levels in the UK Biobank. Luteinizing hormone is from primary care records (Methods), while Testosterone, SHBG, and CBAT are measured or derived from the baseline assessment center visit.*

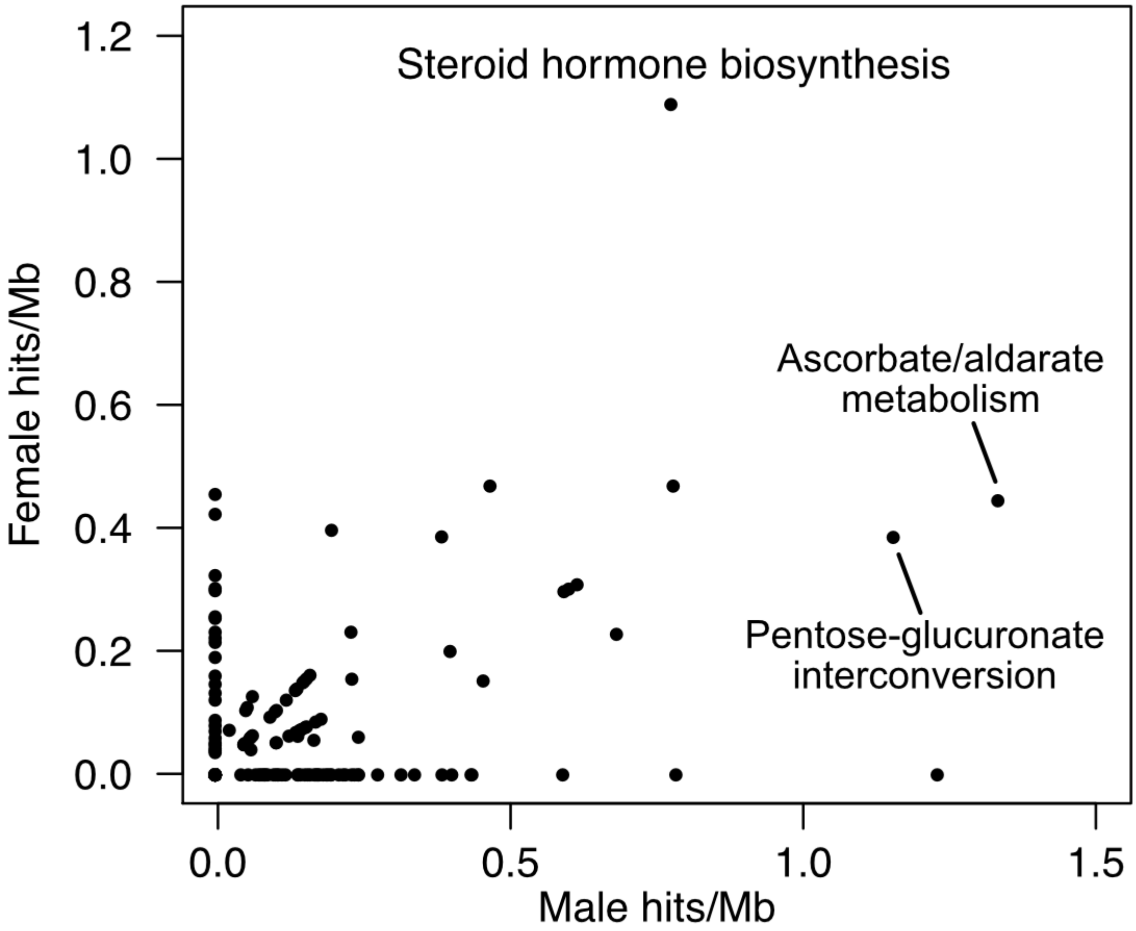


**Figure 6-figure supplement 1:** *The KEGG pathway for steroid hormone biosynthesis is enriched for hits in both female and male testosterone GWAS. The enrichments for male hits in "Ascorbate/aldarate metabolism" and "Pentose-glucuronate interconversion" are almost entirely driven by the UGT genes, which are part of "Steroid hormone biosynthesis"*

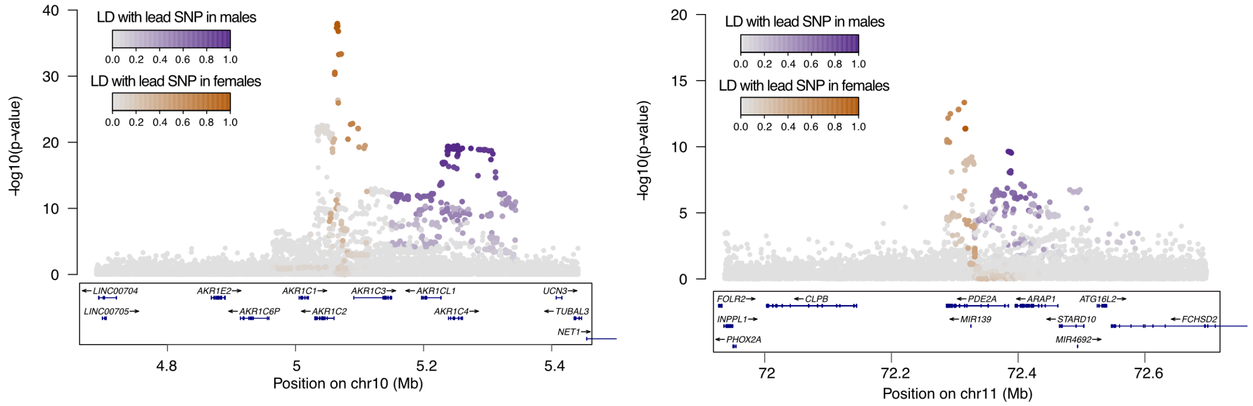


**Figure 6-figure supplement 2:** *Non-overlapping female and male GWAS signals at AKR1C (left) and PDE2A (right) loci.*

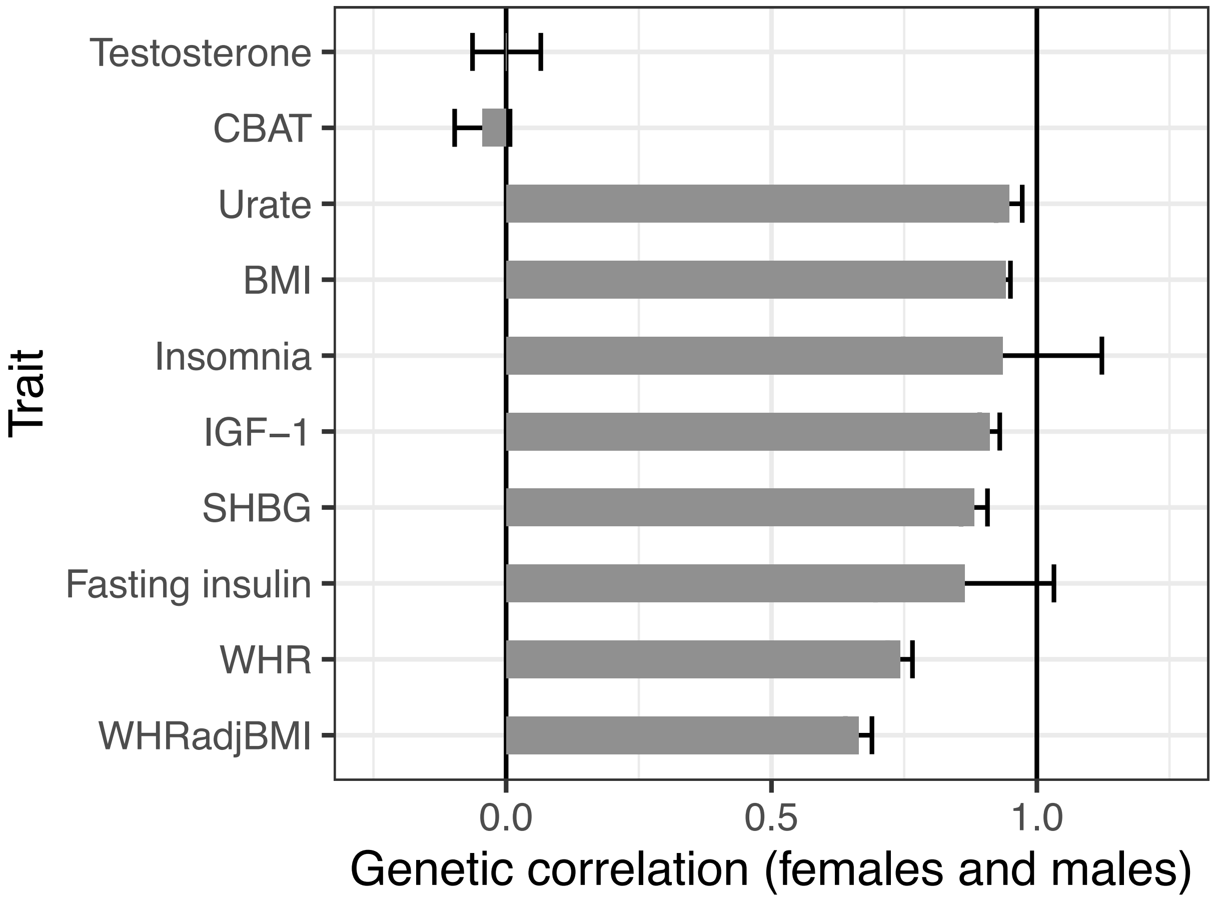


**Figure 7-figure supplement 1:** *Genetic correlations between females and males across select traits. Both Testosterone and calculated bioavailable testosterone (CBAT) show very little genetic correlation, in stark contrast to most other complex traits.*

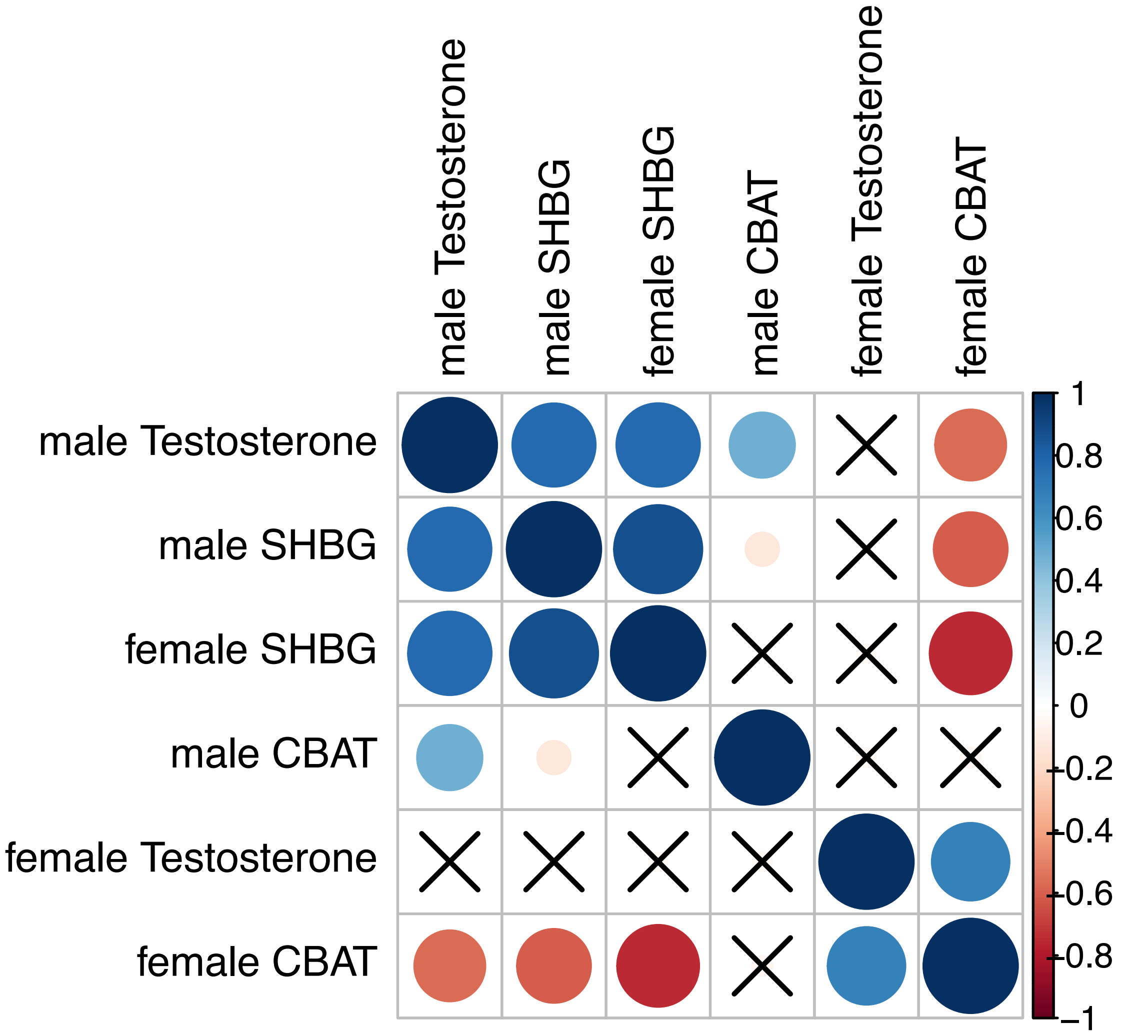


**Figure 7-figure supplement 2:** *Genetic correlations (estimated by LD Score Regression) between total testosterone, SHBG, and calculated bioavailable testosterone (CBAT) in females and males. "X" indicates a non-significant genetic correlation.*

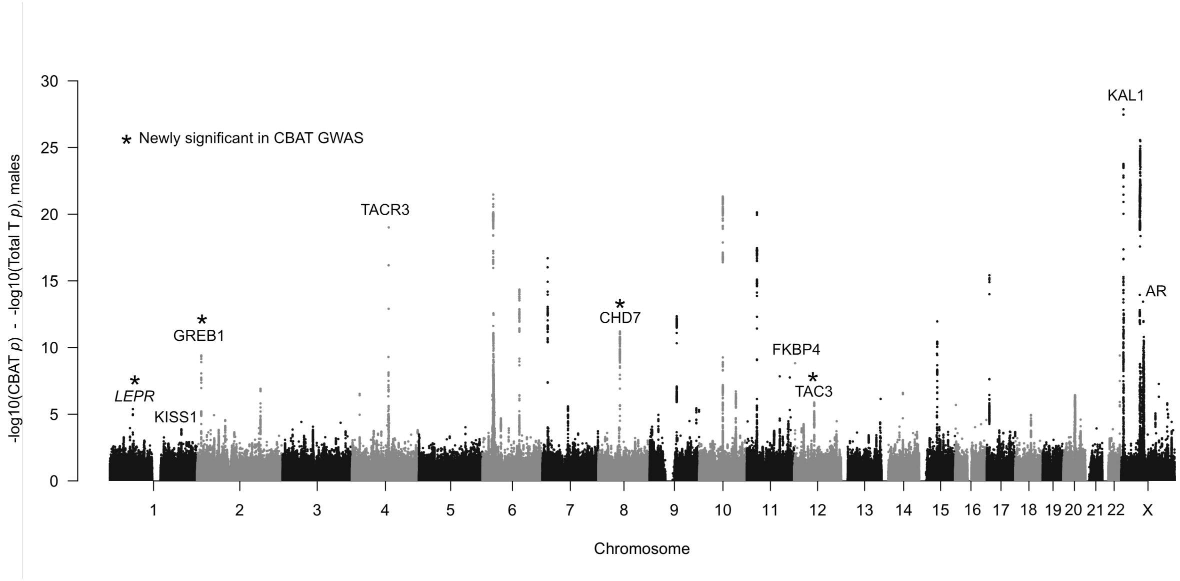


**Figure 7-figure supplement 3:** *Manhattan plot of difference in significance of assocation comparing GWAS of calculated bioavailable testosterone (CBAT) to total testosterone. Only points more significant in CBAT GWAS (i.e. positive values on y-axis) are shown. Genes with known roles in the HPG axis are annotated.*

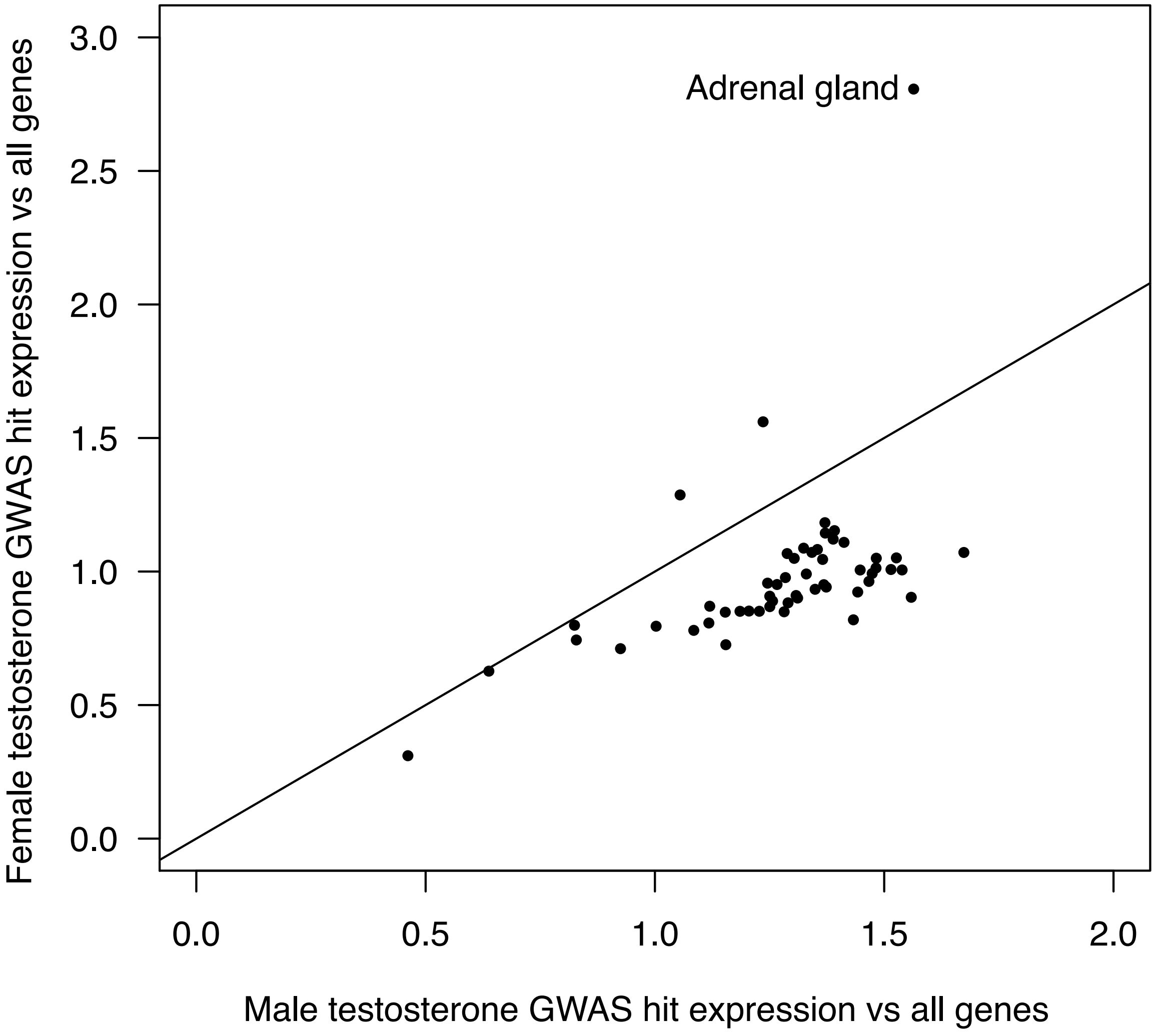


**Figure 7-figure supplement 4:** *Mean expression of testosterone GWAS hits in females or males (defined as mean log-transformed counts of hits divided by mean log-transformed counts of all genes) in each of 48 GTEx tissues.*

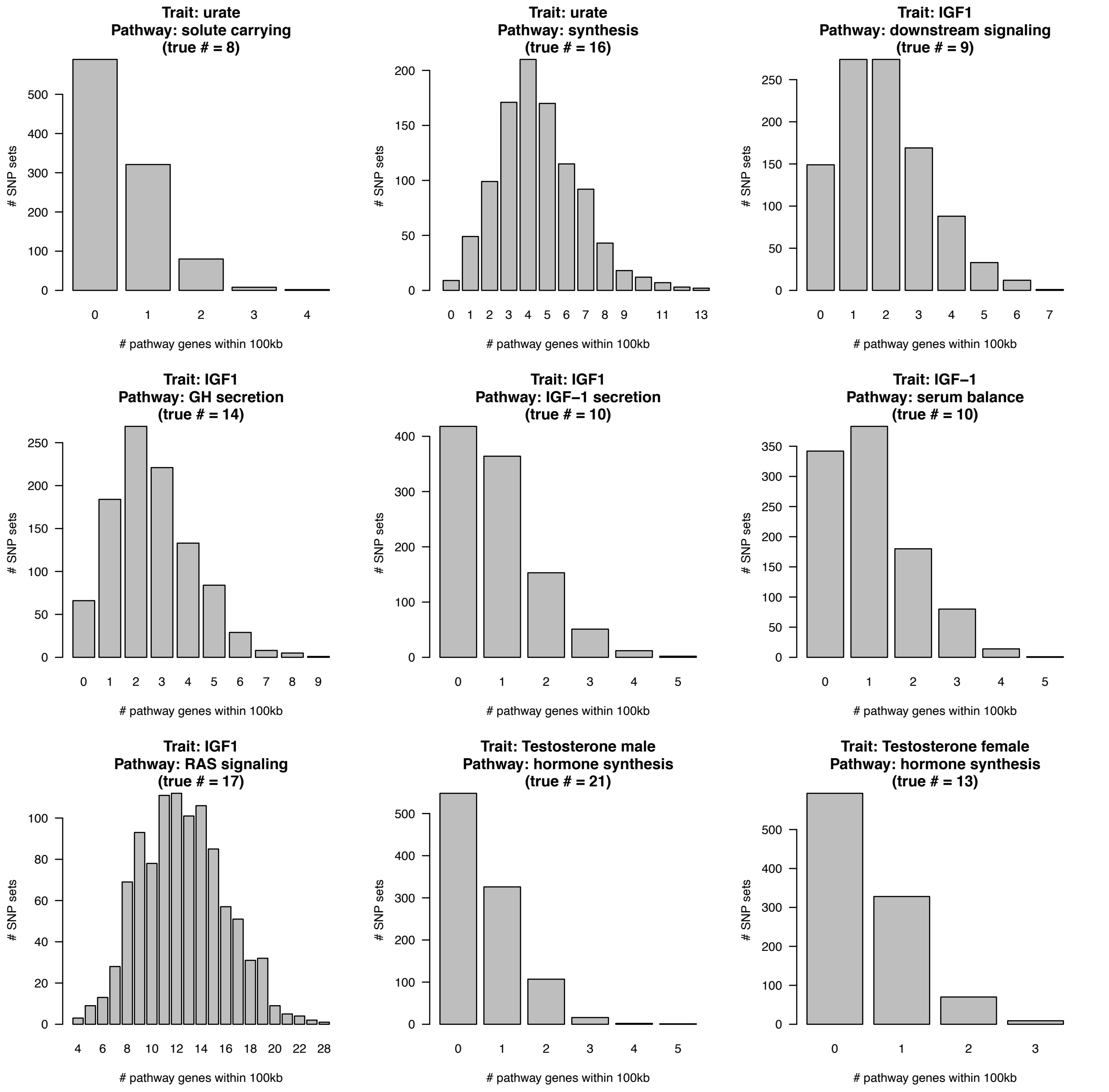


**Figure 7-figure supplement 5: Enrichment of random matched SNPs in core pathways.** *For each GWAS indicated, 1000 sets of equally-sized random SNPs matched to GWAS SNPs for LD, allele frequency, and genic distance (see Methods) were overlapped with the indicated core pathway with 100kb windows. The*

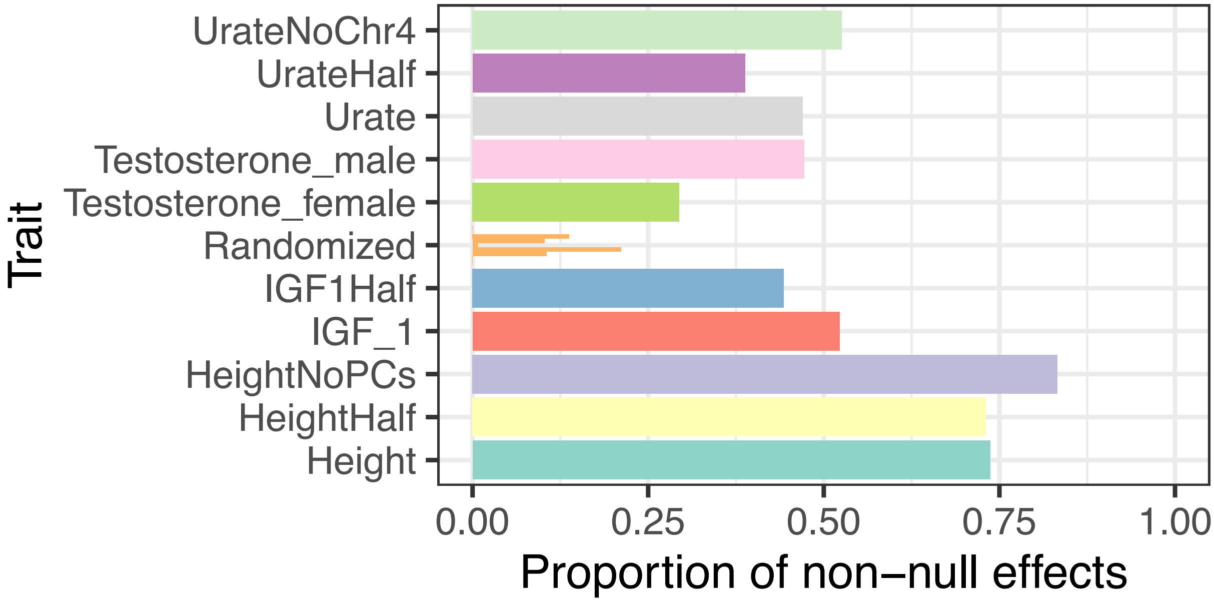


**Figure 8-figure supplement 1:** *Proportion of non-null associations in a random sample of 100,000 variants for each trait. Variants are then filtered to those with minor allele frequency greater than 5%, and ashR is run on the beta and standard error to estimate the proportion of non-null associations. "UrateNoChr4" is the Urate GWAS with chromosome 4 excluded, where the two largest effect loci, SLC2A9 and ABCG2, are both located. Traits with the suffix "Half" are 50% downsamplings of the White British cohort to approximate the sample sizes of the sex-stratified testosterone GWAS. "Randomized" are random associations generated by evaluating shuffled versions of the Urate (n = 3 shuffles) and IGF-1 (n = 3 shuffles) phenotypes and represent estimates we might expect under the null distribution of no associations, where we observe some noise but consistently observe estimates well below that of un-shuffled traits. These global estimates represent the proportion of all variants in a given trait that are linked to causal sites.*

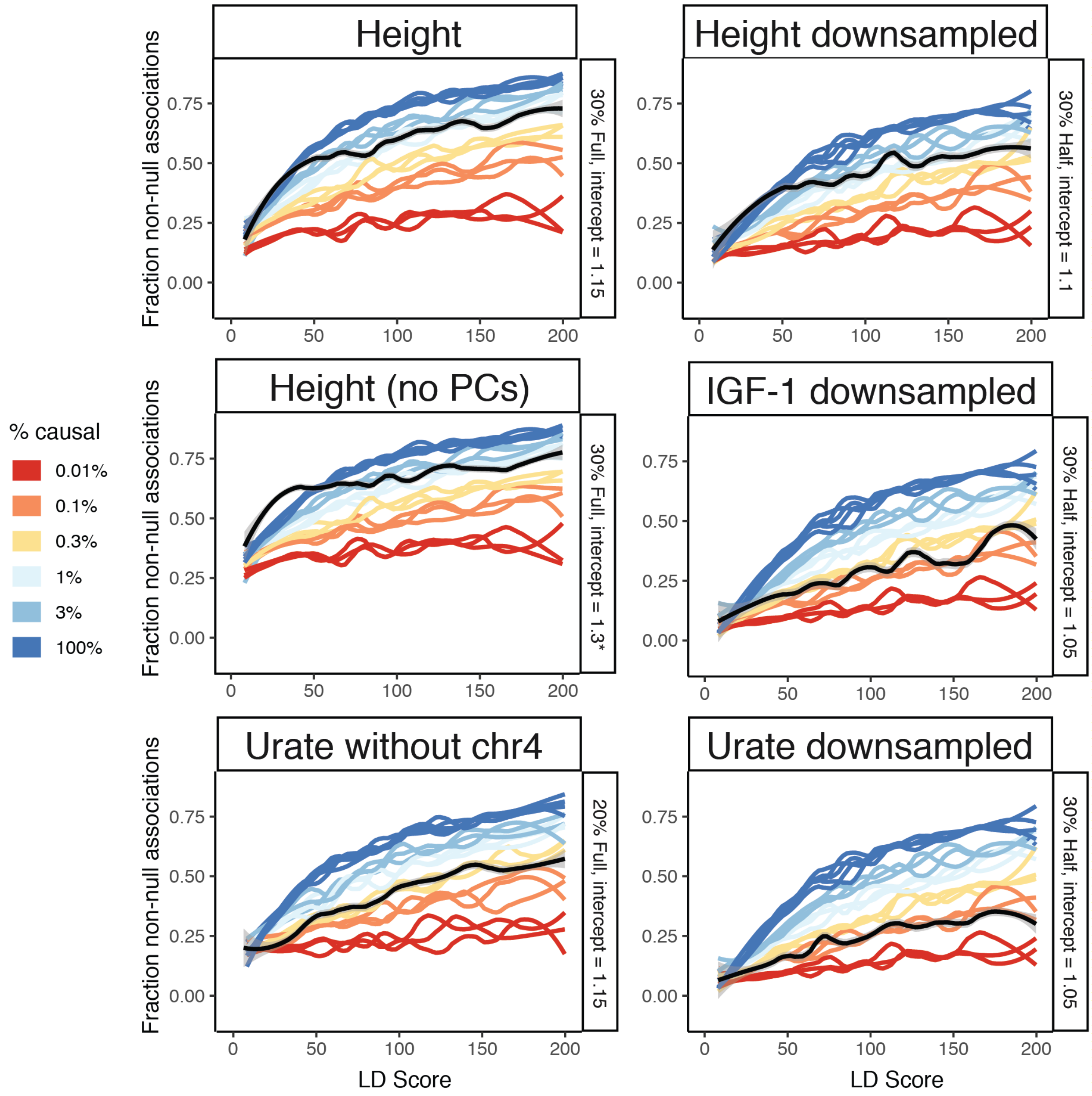


**Figure 8-figure supplement 2:** *Additional traits to fit causal simulations on. Height, height without principal components regressed out of the GWAS, Urate with chromosome 4 excluded (where both SLC2A9 and ABCG2 are located, reducing SNP-based heritability to ≈ 20%), and random 50% downsamplings of height, IGF-1, and urate (including chromosome 4) are shown. See Figure 8-figure supplement 3 for additional sex hormone traits. Despite matching simulations for sample size, slight reductions in causal variant count are still observed in downsampled traits, suggesting that power is still possibly limiting, of particular relevance to the sex-stratified testosterone GWAS. As expected, excluding principal components from the GWAS dramatically increases the intercept. * 1.3 was the highest tested intercept and is still too low, but no more runs were completed as the curve is clearly shaped very differently than simulation runs regardless of intercept. We recommend performing GWAS while regressing out principal components to avoid difficult to interpret results. See Figure 8-figure supplement 3 for additional fits to sex hormone related traits.*

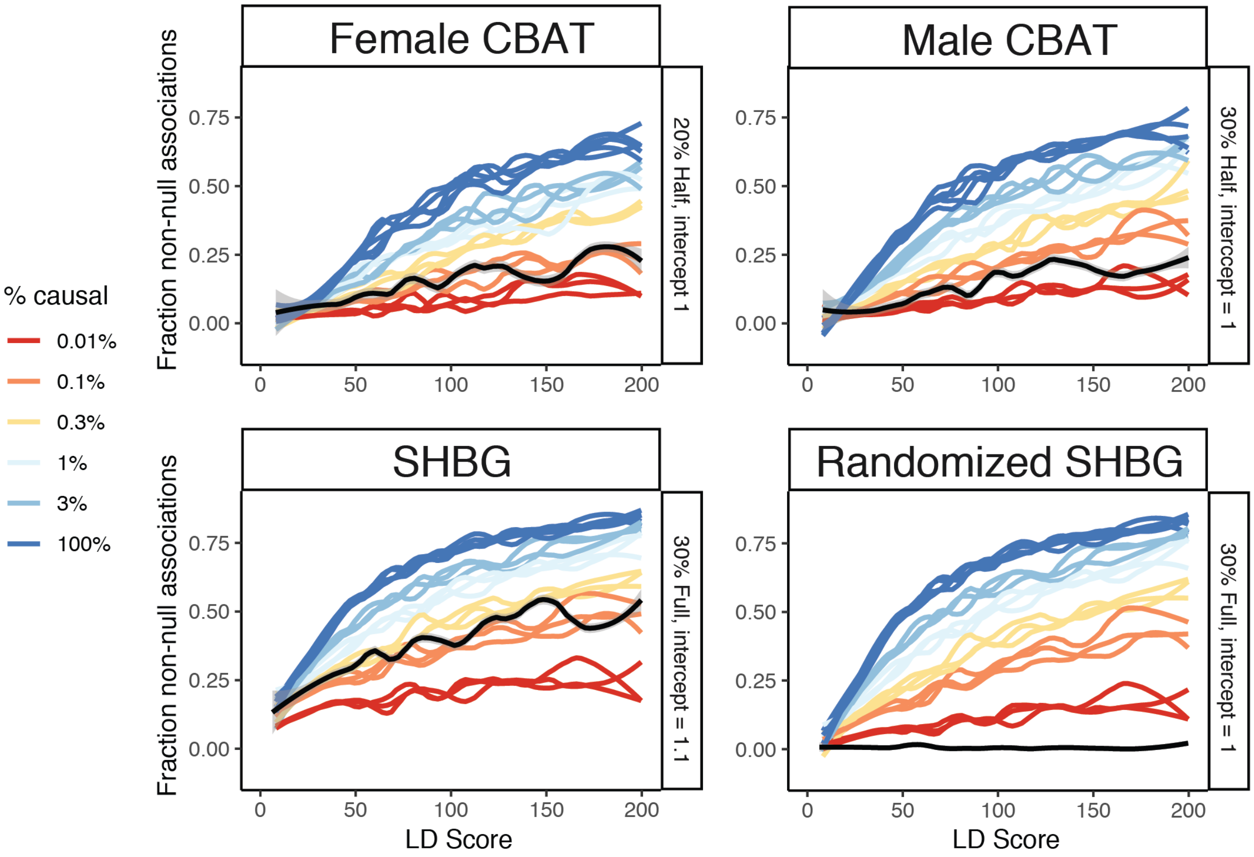


**Figure 8-figure supplement 3:** *Prediction plots for the causal SNP counts underlying calculated bioavailable testosterone (CBAT) in females and males, as well as sex hormone binding globulin (SHBG) and a randomized version of SHBG. CBAT traits are similar to their testosterone counterparts and SHBG is estimated to have slightly more causal variants (≈ 0.2%)*

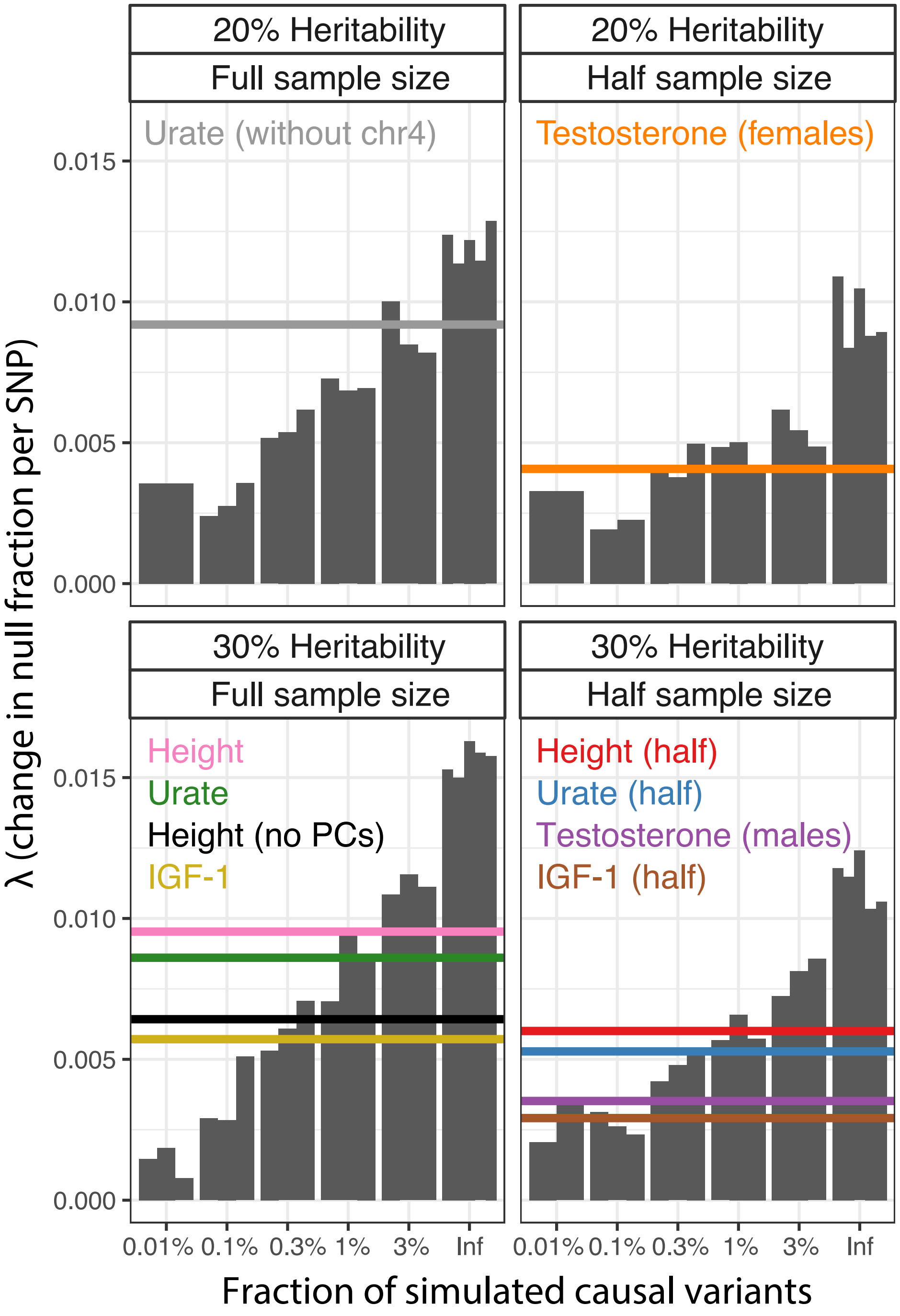


**Figure 8-figure supplement 4:** *Parametric causal fraction on LD Scores reproduces SNP-based heritability-based estimates. For each study sample size (Full, full UK Biobank White British; Half, 50% downsample [and sex-stratified]) and SNP-based heritability (20% or 30%), simulations run with different causal variant proportions ($n = 3$ reps except infinitesimal, $n = 5$) were used as a background to estimate the causal site proportions for each complex trait. Complex trait non-linear least squares $\lambda$ estimates are shown as horizontal lines.*

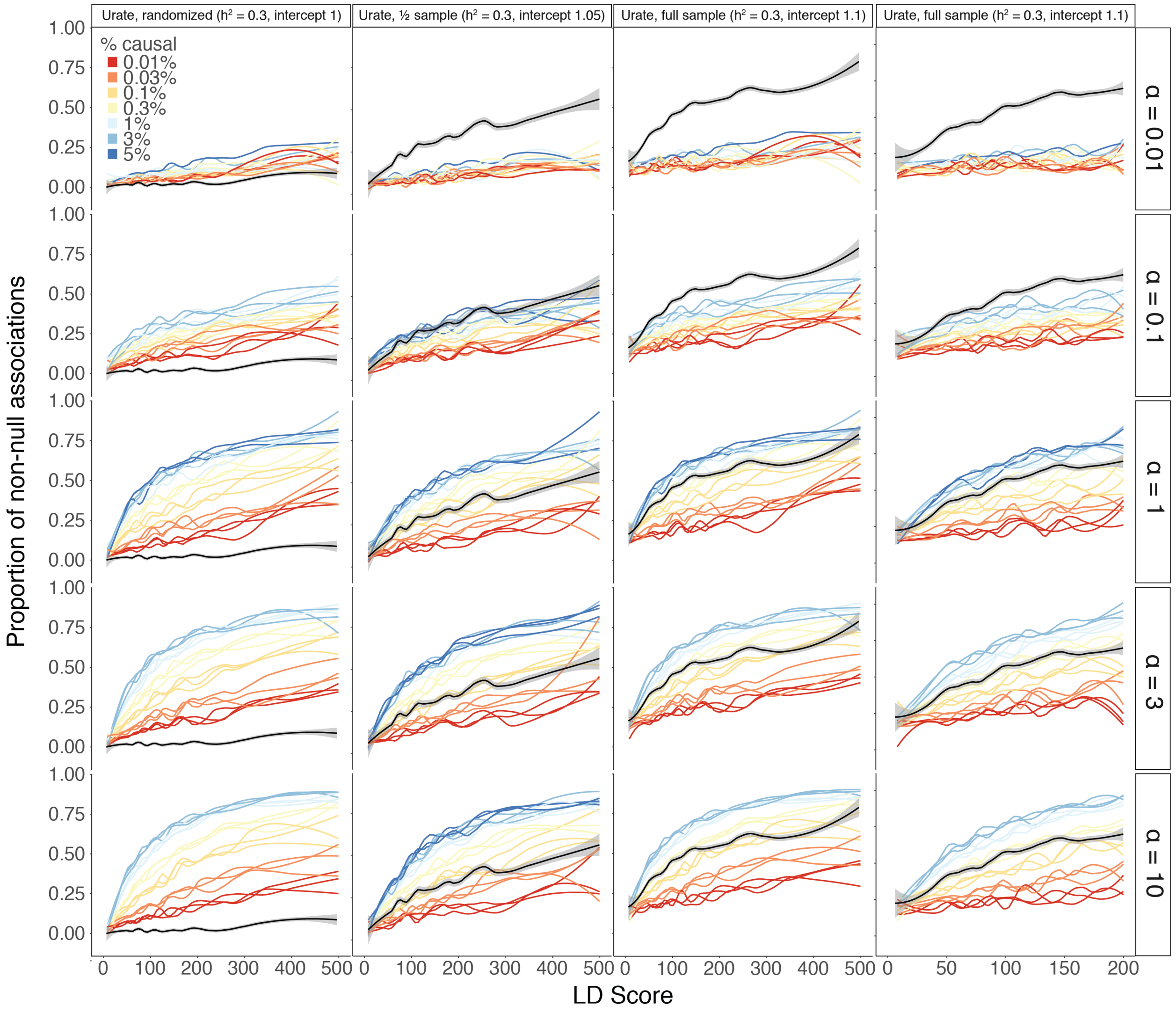


**Figure 8-figure supplement 5:** *Estimates of causal sites are conservative with respect to SNP concentration within the genome. Rather than drawing SNPs uniformly at random from those with MAF > 1% (Methods), instead each megabase window was assigned a distinct probability $\rho$ of SNPs being causal within that megabase window. For a given causal fraction c, $\rho$ was drawn at random from $Beta(\alpha, \alpha/c)$. Thus, the mean causal fraction across all megabase windows in the genome is still c, and this fraction is concentrated more in single windows under decreasing values of $\alpha$. The standard results shown in main text correspond to $\alpha = \infty$. As we decrease $\alpha$, the estimates consistently increases, regardless of the same size or LD Score axis threshold. However, the results consistently stay above the randomized version of the trait, suggesting that the estimates remain non-zero even for very concentrated SNP-based heritability.*

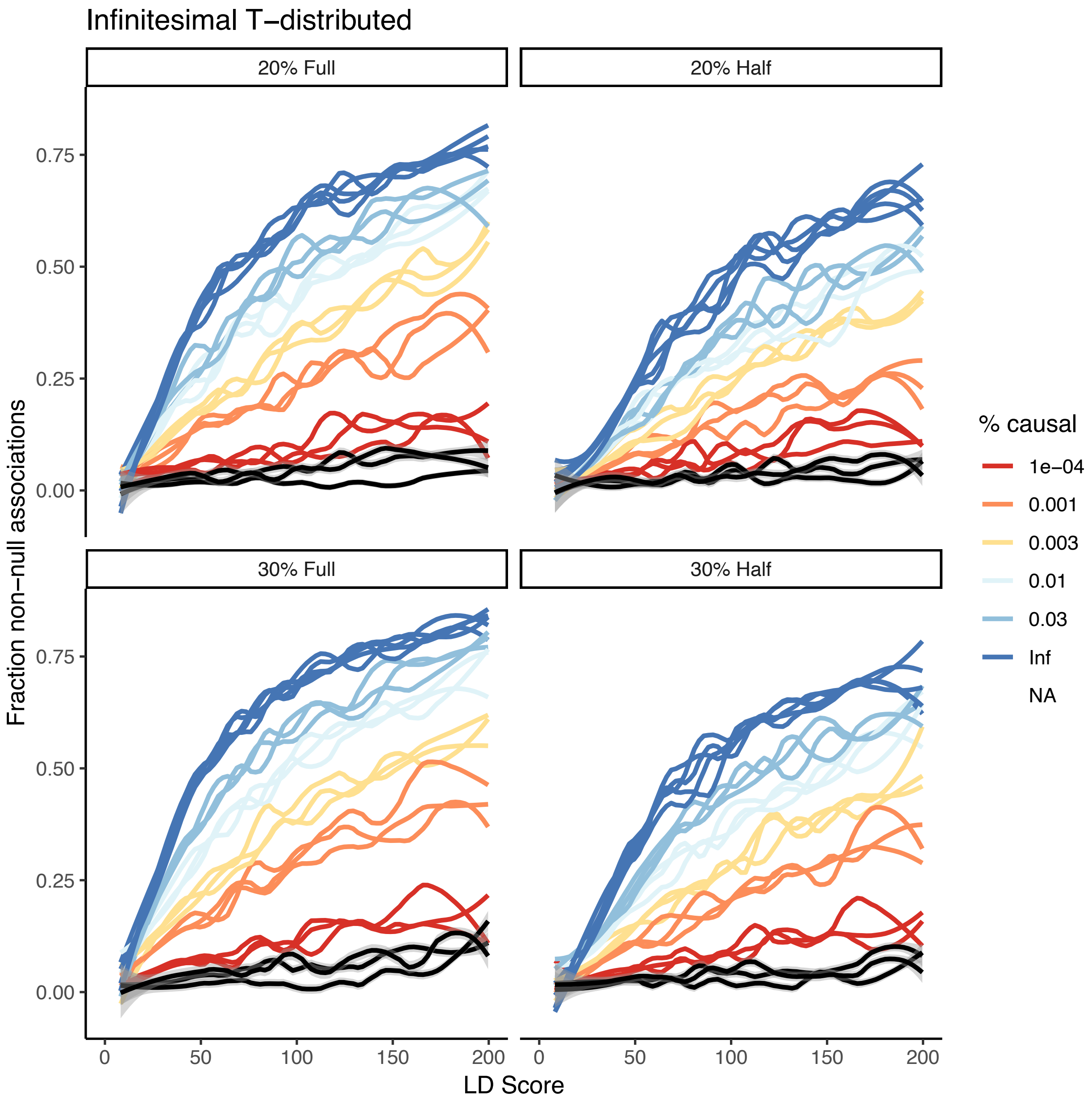


**Figure 8-figure supplement 6:** *Effect of distribution of causal site betas on estimates of causal variant count. A model where every variant in the genome is drawn from a one-degree-of-freedom T-distribution is shown in black across the different heritabilities and samples sizes. Because some small number of variants have very large effects in the T-distribution, and the total SNP-based heritability is normalized to this additional variance, the estimate is downward biased and suggests that rather than 100% of variants being causal, instead less than 0.01% are causal. In practice, most effect sizes are not as overdispersed as a T-distribution, though some traits (e.g. Urate) have a few variants with very large effect.*

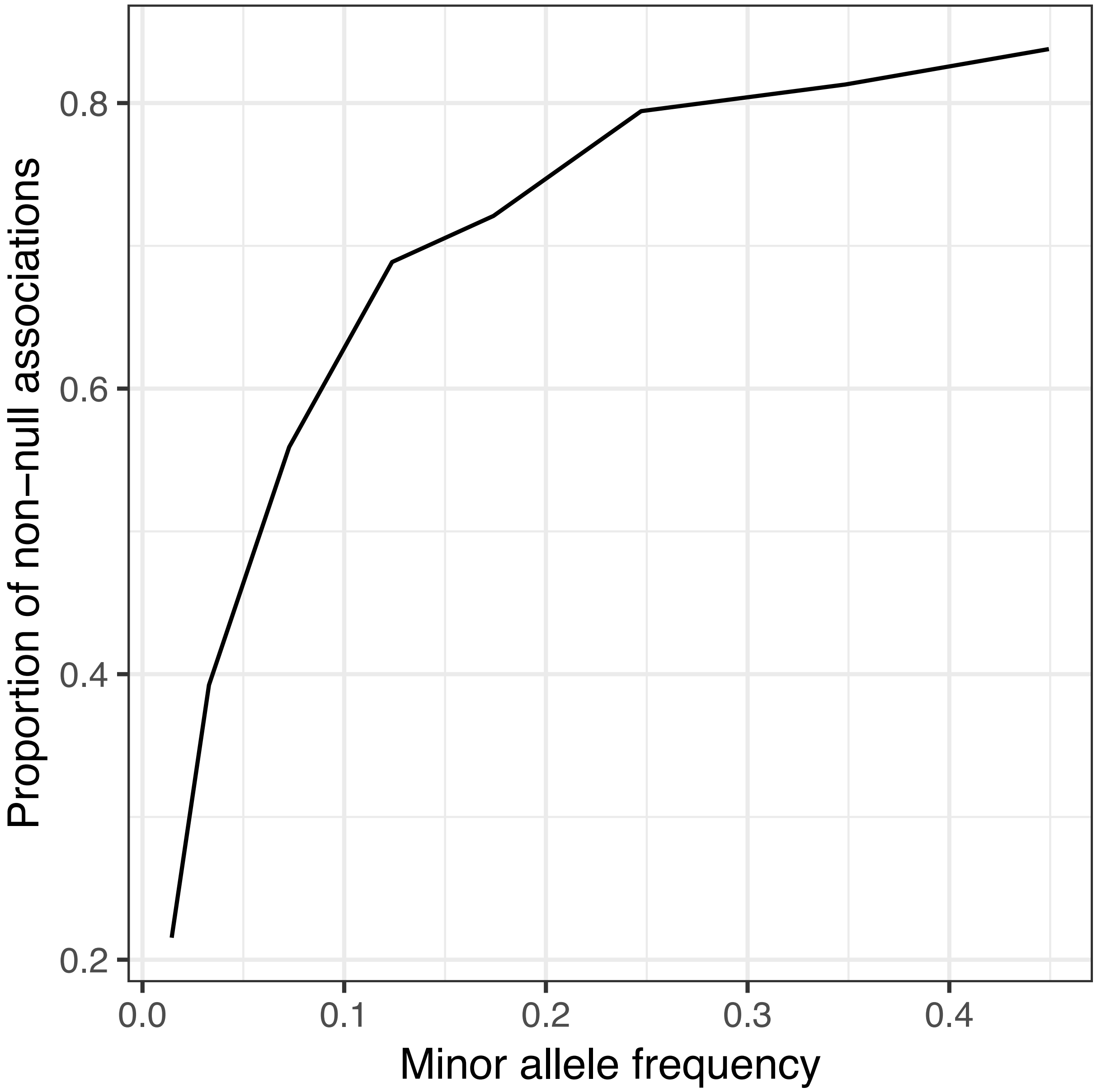


**Figure 8-figure supplement 7:** *Association between minor allele frequency and estimated proportion of causal variants. Simulated traits were generated with every variant with MAF > 1% being causal (with effect size drawn from N(0,1) independently) and a GWAS was performed with* $h^2 = 0.3$*. Then, for each MAF bin, 100,000 variants were sampled at random, and ashR was run to estimate the proportion of causal variants (or variants in LD with a causal variant) within each bin. There is a strong relationship with minor allele frequency, suggesting that power might still be a consideration even at UK Biobank sample sizes. See also Figure 8-figure supplement 8.*

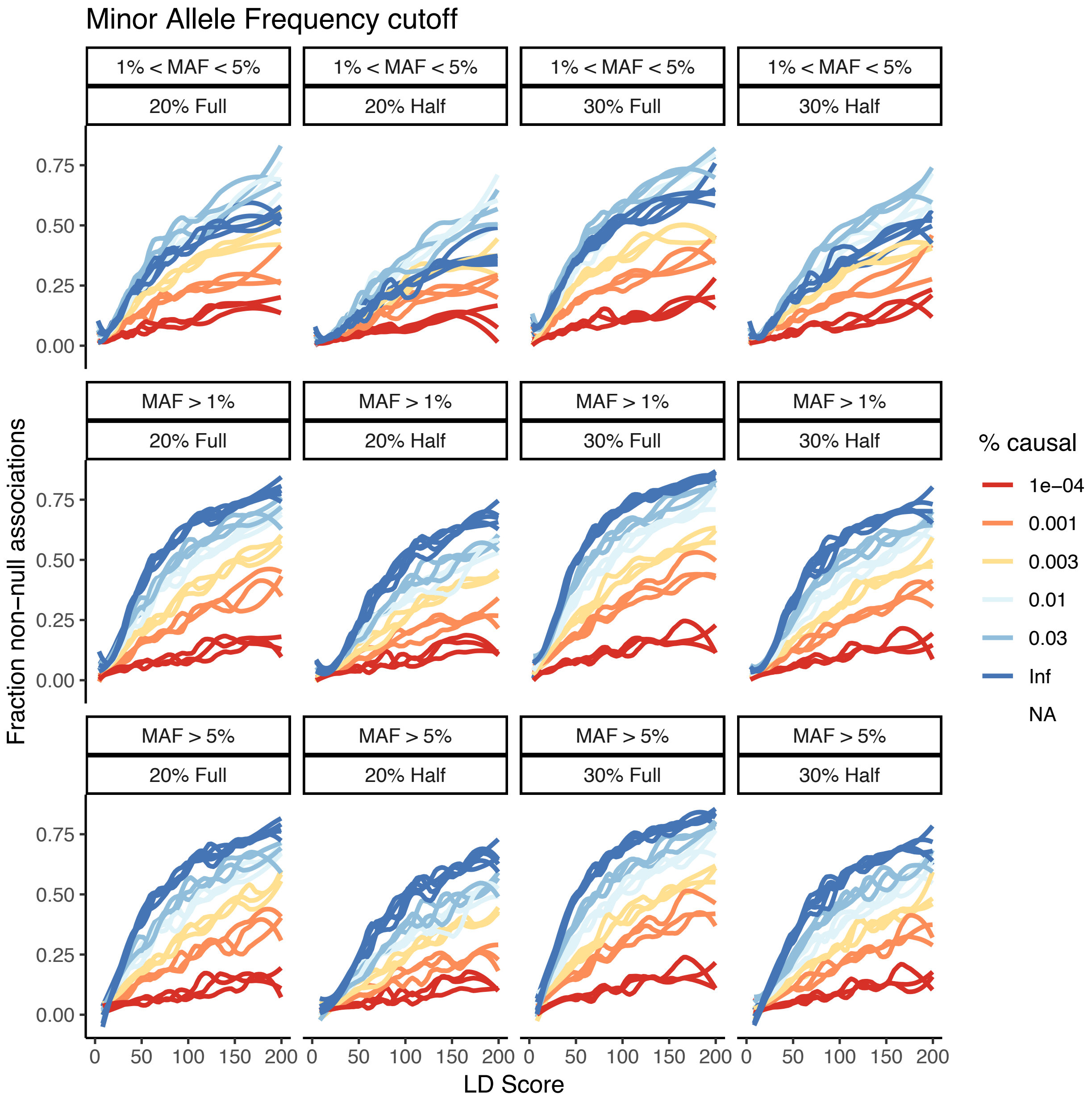


**Figure 8-figure supplement 8:** *Effect of minor allele frequency cutoff on the estimates obtained. Three choices are considered – MAF > 1%, MAF > 5%, and MAF between 1% and 5%. There is some attenuation of the estimates in the MAF between 1% and 5% analysis, and the infinitesimal estimates are below that of the models with fewer causal sites, supporting the notion that power is still limiting.*

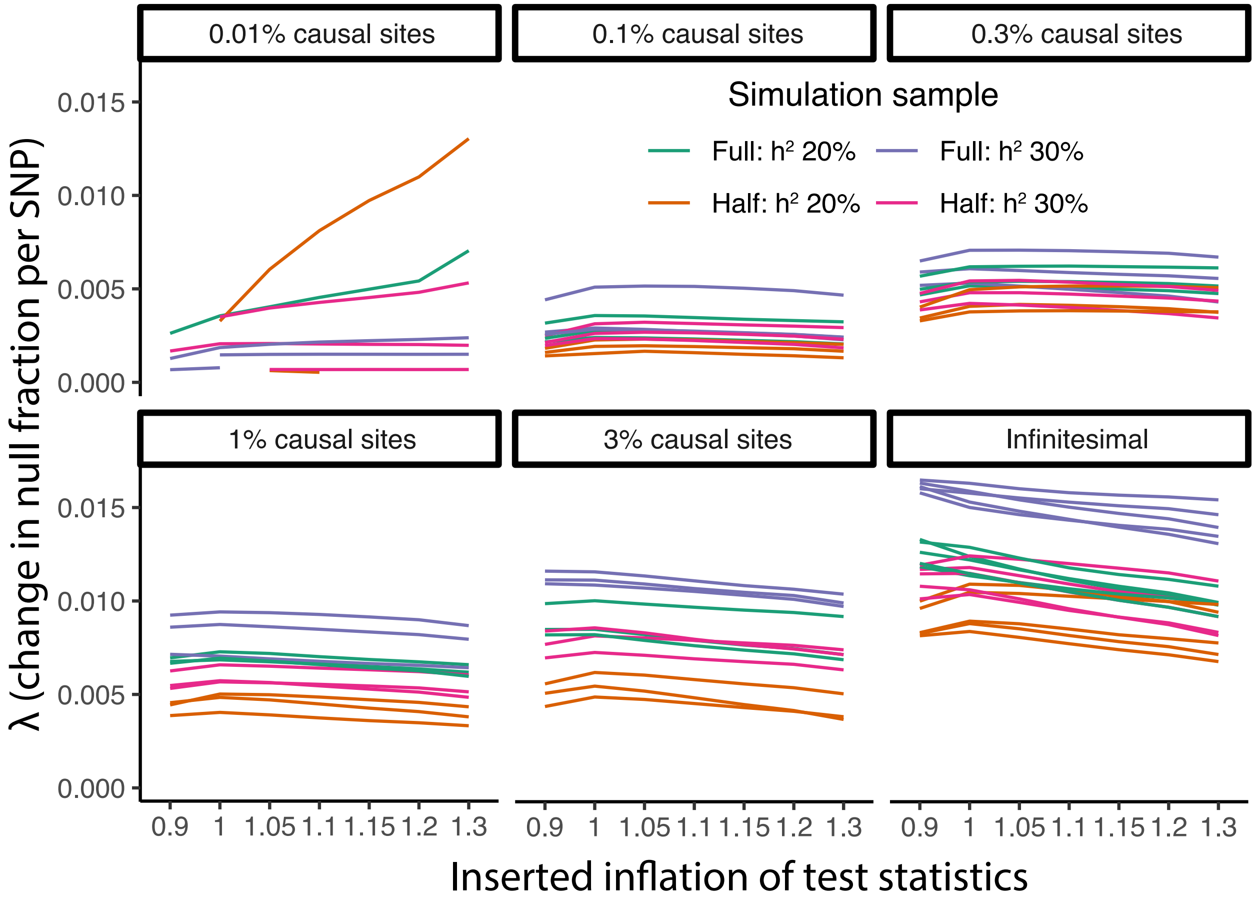


**Figure 8-figure supplement 9:** *Parametric causal fraction is robust to population structure. Within each causal site proportion (faceted), inflation added by systematically reducing standard errors was applied with ratios 0.9 through 1.3. For each, the parametric λ fit was calculated using non-linear least squares (Methods). Across this range of inflation levels, there was no large or consistent change in λ estimates, suggesting the λ fit is relatively robust to population structure.*

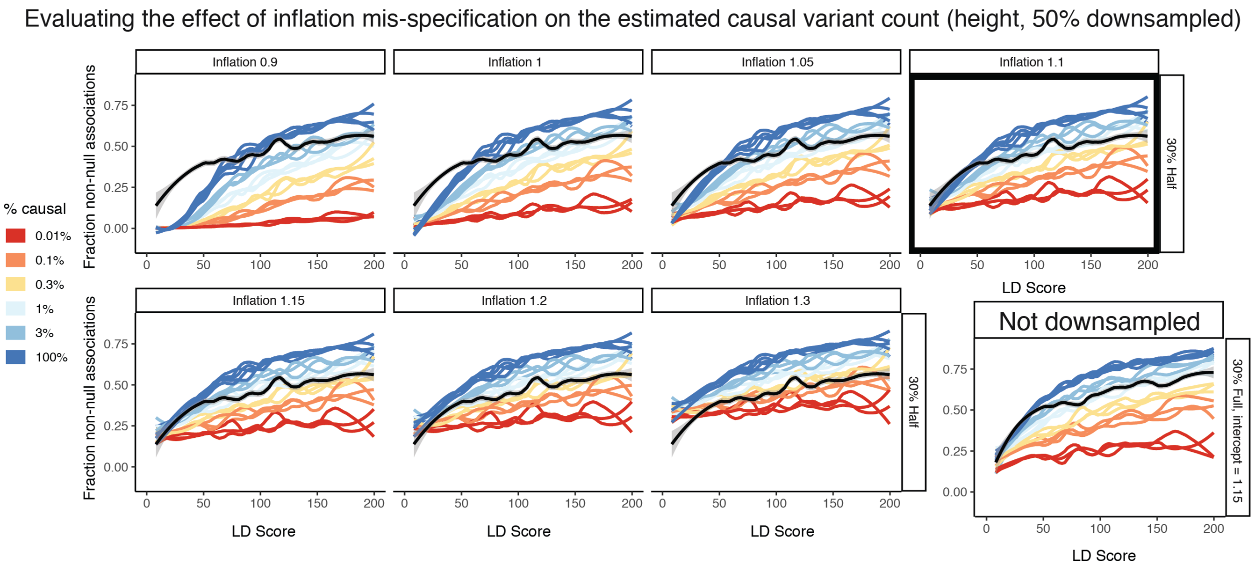


**Figure 8-figure supplement 10:** *Estimating the effect of inflation mis-specification on the estimated causal variant count. 50% downsampled height is used as an example. When changing the inflation, dramatic differences in the best fitting curve are possible in the simulation matching approach, ranging from greater than 3% at an inflation of 0.9 to around 0.2% for an inflation of 1.3. This is in part exacerbated by the reduced difference between the simulations with different causal percentages at high inflation levels. The best intercept is chosen on the basis of the closest fit at an LD Score of 0 (selected in black box, intercept = 1.1). Note that sample size can change this intercept for the same trait, as shown by the full sample height results (where best intercept = 1.15). However, the resulting estimate is the very similar conditional on intercept.*

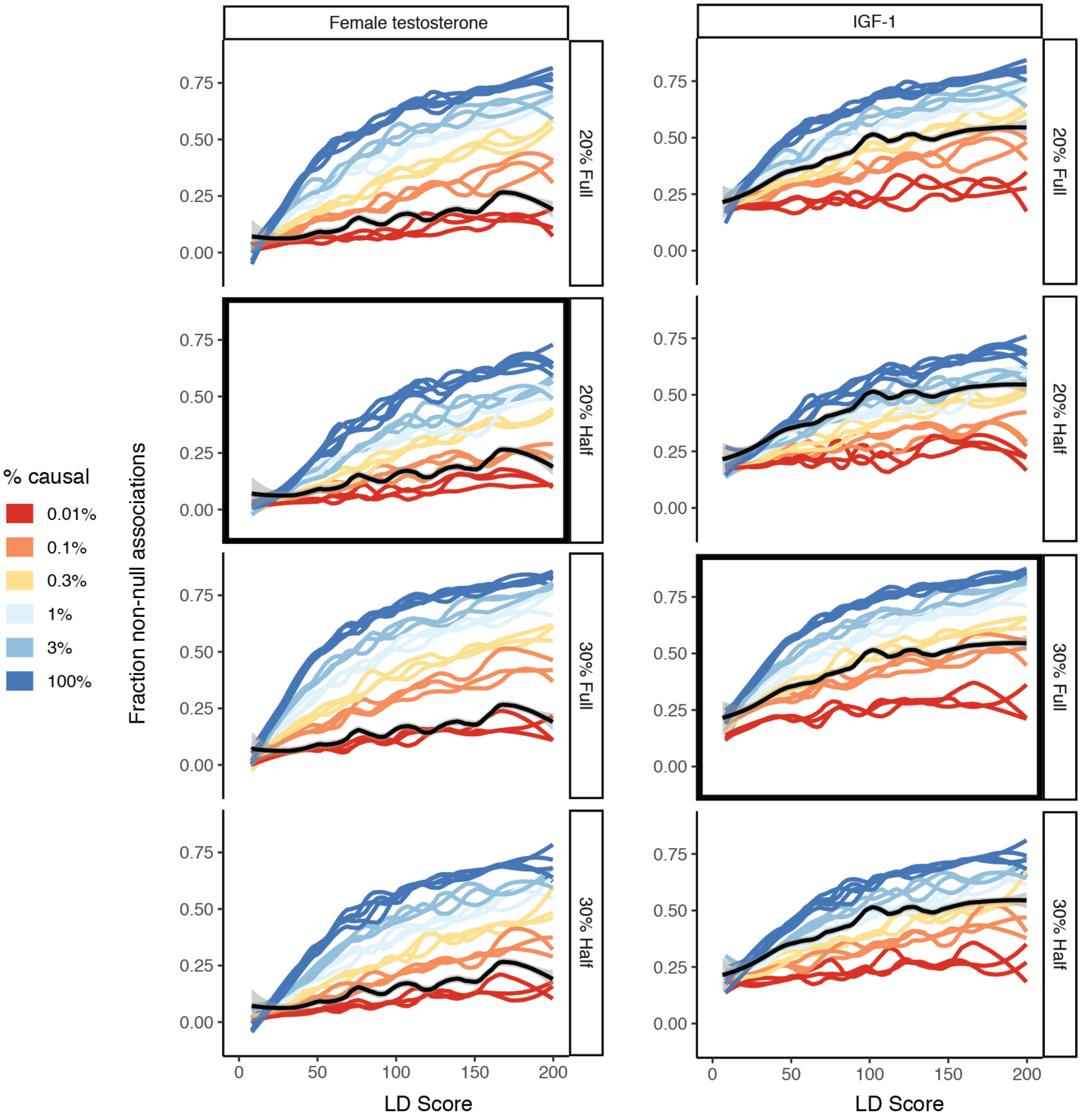


**Figure 8-figure supplement 11:** *Effect of mis-specification of SNP-based heritability or sample size in the simulation matching approach. For female testosterone (left) and IGF-1 (right), the correct specification that corresponds to the trait is selected in black. All other estimates are given as well, revealing that the choice of SNP-based heritability and sample size does have a moderate effect on the resulting estimates. As such, we recommend matching the sample size and SNP-based heritability of the simulation GWAS as closely as possible to the traits under study.*

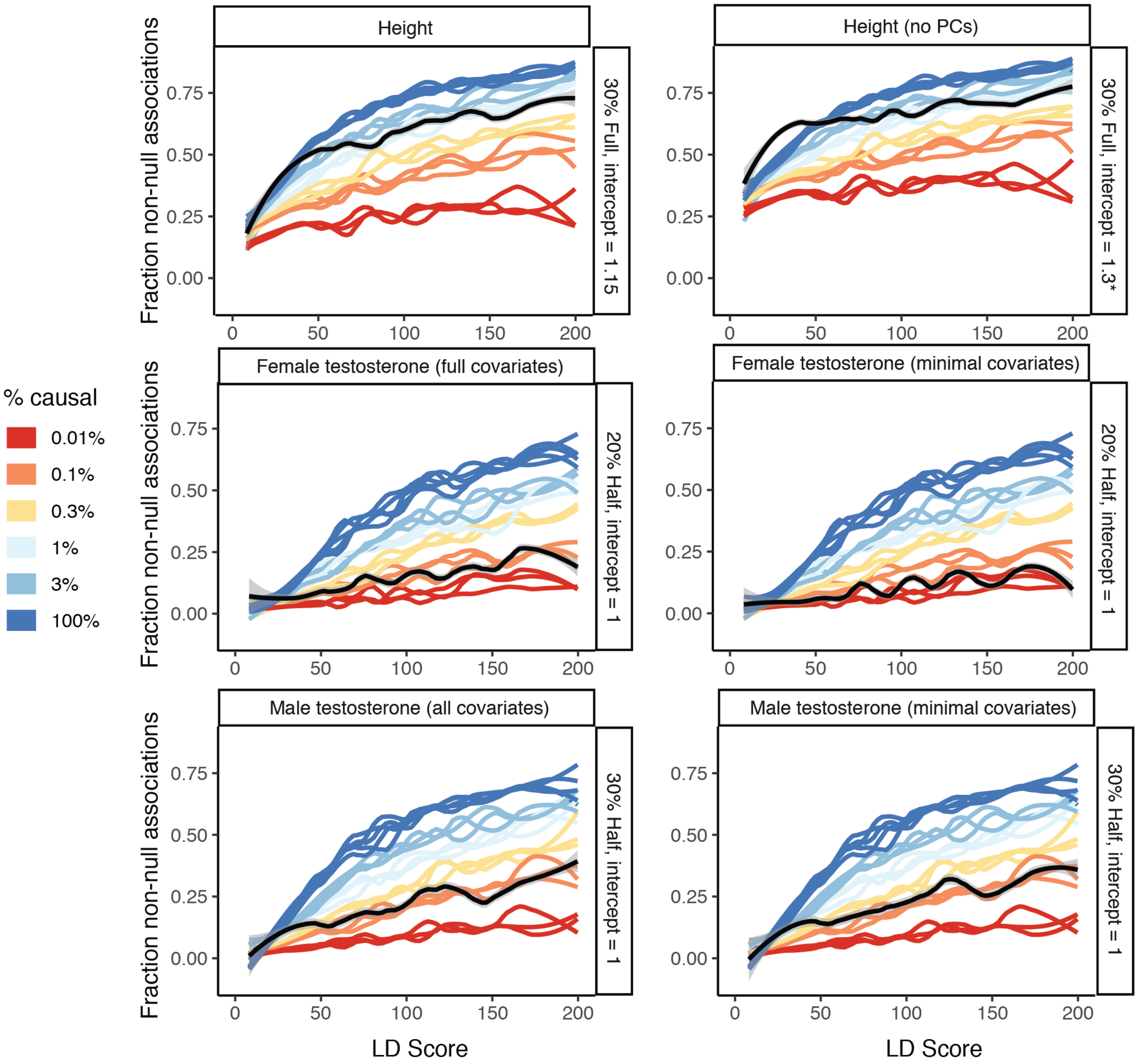


**Figure 8-figure supplement 12:** *Effect of GWAS covariates on estimates. As noted in Figure 8-figure supplement 2, not including principal components can meaningfully effect estimates and produce upward bias, so we recommend adjusting for principal components (or using a suitable mixed model). In addition, we evaluated testosterone traits using two covariate sets: the complete set and a minimal age, age$^2$, genotyping array, and principal components 1-20 that were used for the CBAT GWAS (due to the derivation from albumin, SHBG, and testosterone, and the difference in batch between these, Methods). For male testosterone the effect of different covariates was negligible, while for female testosterone, the estimate was slightly lower with the minimal covariates. We recommend optimizing covariates to the sample and traits under study to maximize the interpretability of the causal variant estimation.*

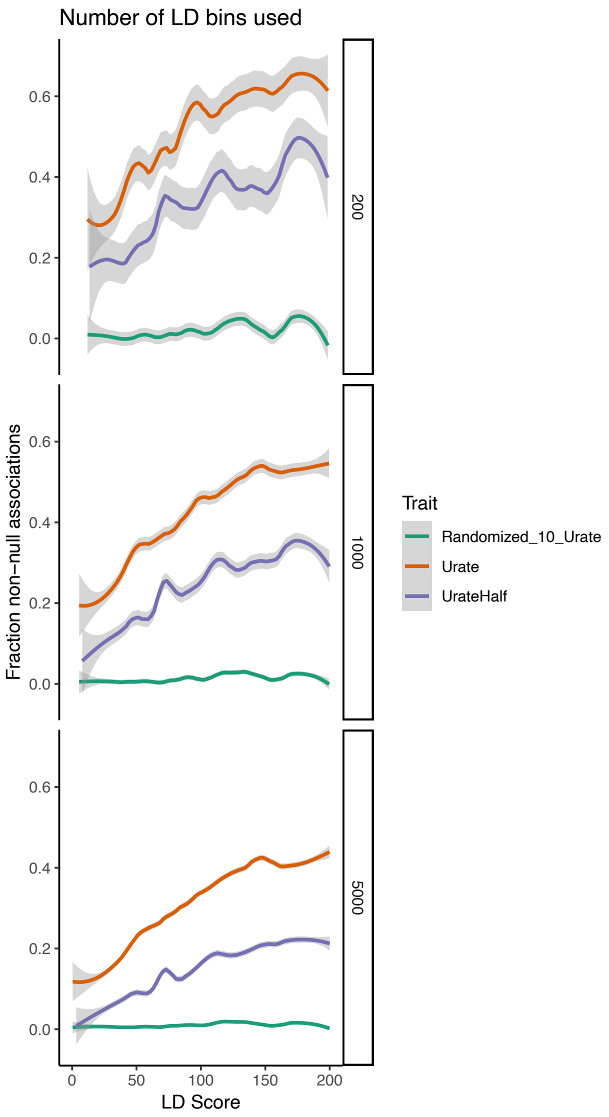


**Figure 8-figure supplement 13:** *Effect of bin count on estimates of causal variants. For each binning option, urate, randomized urate, and urate 50% downsampled were run. 1000 bins, 5000 bins, and 200 bins were compared. Lower intercepts and smaller standard errors were observed with higher bin count. We recommend that as many bins as is practicable are used for these analyses.*

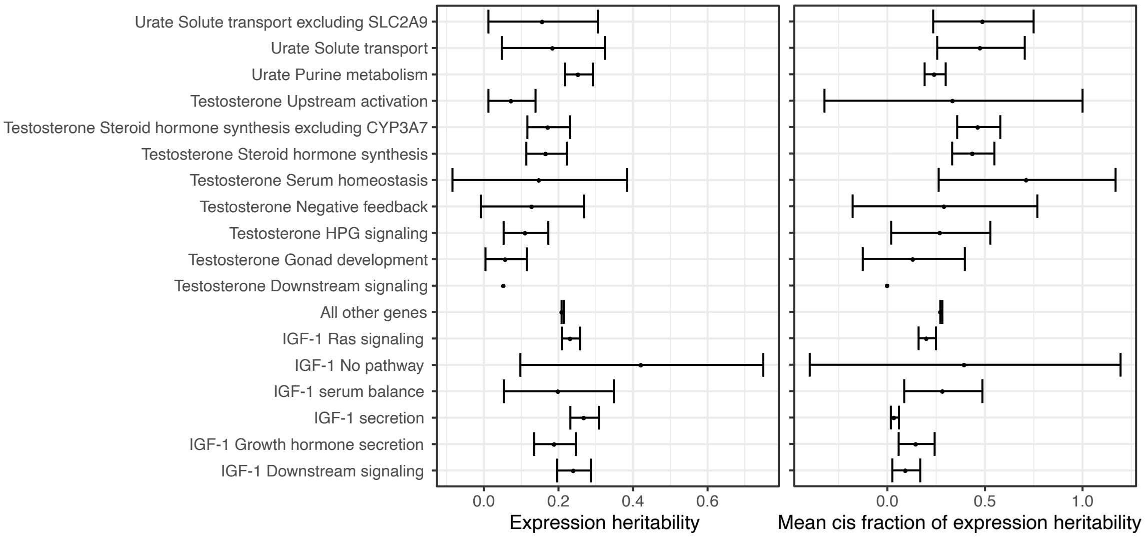


**Figure 8-figure supplement 14:** *Distributions of (left) total SNP-based heritability of gene expression, or (right) fraction of expression SNP-based heritability driven by cis-effects [126] for genes in the indicated core pathways, or for all other MsigDB genes not in a core pathway. Points indicate means, error bars indicate two standard deviations*
